# Microglia-astrocyte interaction underlies aquaporin-4 dysregulation in the mouse cortex

**DOI:** 10.64898/2025.12.03.691996

**Authors:** Yan Wang, Emiko Morita, Yuka Hirota, Hiromu Monai

**Author notes:** To whom correspondence should be addressed: Hiromu Monai.

## Abstract

Brain water homeostasis relies on the coordinated actions of glial cells. In particular, astrocytes play an essential role in facilitating perivascular water exchange between cerebrospinal fluid and interstitial fluid through aquaporin-4 (AQP4), a water channel highly concentrated at astrocytic endfeet. Dysregulation of AQP4 localization is implicated in various neuropathologies, but its underlying mechanisms are unclear. Although astrocytes and microglia both express β-adrenergic receptors (β-AdRs), whether β-AdR signaling modulates astrocytic AQP4 polarization through microglial activation has not yet been investigated. Here, we hypothesized that microglial activation mediates the β-AdR-induced loss of astrocytic AQP4 polarization. To test this hypothesis, we topically applied the β-AdR agonist, isoproterenol, to the primary visual cortex of anesthetized mice and evaluated AQP4 polarization using double immunohistochemistry. Acute β-AdR activation (3 h) significantly reduced perivascular AQP4 polarization and enhanced local microglial reactivity. Both pharmacological inhibition of microglial activity with minocycline and microglial depletion via dietary administration of the CSF1R antagonist, PLX5622, prevented AQP4 dysregulation induced by isoproterenol. These findings demonstrate that microglial activation is required for β-AdR agonist-induced AQP4 dysregulation in the mouse cortex, revealing a previously unrecognized microglia-astrocyte interaction linking adrenergic signaling to glial water homeostasis.

## Introduction

Aquaporin-4 (AQP4), a crucial member of the aquaporin family, is predominantly expressed in the central nervous system, particularly in the foot processes of astrocytes surrounding capillaries, beneath the meninges, and within the ventricles[1,2]. AQP4 not only plays a vital role in maintaining water and potassium homeostasis in the brain [3–5], but also participates in regulating extracellular space volume and modulating neuroinflammation [2]. In particular, AQP4 polarization at astrocytic endfeet facilitates efficient cerebrospinal fluid-interstitial fluid exchange through the glymphatic system, contributing significantly to brain clearance [6–8].

As a water channel protein, AQP4 plays a crucial role in the development and recovery of cerebral edema [9,10]. However, AQP4 depolarization exerts differential effects on various types of brain edema [11]. While AQP4 deficiency aids in the recovery from cytotoxic edema in mice [12], it is detrimental in cases of vasogenic edema [13]. Frydenlund et al. have found that in mice with transient cerebral ischemia induced by middle cerebral artery occlusion (MCAO), polarized AQP4 decreased 24 hours after reperfusion, with subsequent recovery of AQP4 expression at 72 hours post-reperfusion contributing to brain edema resolution [14]. Additionally, Monai et al. have suggested that pan-adrenergic receptor (AdR) antagonists may facilitate recovery from acute ischemic stroke by preserving AQP4 expression and polarization in mice [15]. Nevertheless, the precise mechanisms by which AdR signaling activation affects AQP4 polarization remain unclear.

Microglia, immune cells in the central nervous system, are rapidly activated following various neurological diseases, undergoing significant changes in morphology and gene expression. They not only release pro-inflammatory cytokines such as tumor necrosis factor-α (TNF-α), interleukin-1β (IL-1β), interleukin-6 (IL-6), and various chemokines, thereby inducing neuroinflammatory responses, but also participate in pathological processes such as injury spread, alteration of blood-brain barrier permeability, and apoptosis [16–18].

Following acute ischemic stroke, brain tissue rapidly develops energy metabolism disorders and hypoxia due to interrupted blood flow, leading to ion homeostasis imbalance, oxidative stress, and cellular necrosis [19,20]. Under these conditions, microglia undergo rapid morphological transformations, shifting from a resting, slender, multi-processed state to a spherical structure characterized by shortened pseudopodia and enlarged cell bodies, accompanied by significant alterations in gene expression profiles [16,17,21]. Activated microglia massively release pro-inflammatory cytokines (e.g., TNF-α, IL-1β, IL-6) and chemokines (e.g., MCP-1, CXCL10), and upregulate degradative enzymes like matrix metalloproteinase-9 (MMP-9) [22,23]. MMP-9 can disrupt AQP4 polarization by degrading anchoring molecules at the endfeet of astrocytes, leading to an imbalance in water transport between the perivascular space and brain parenchyma, thereby exacerbating vasogenic edema formation [10,24].

Recent studies have demonstrated that microglia express functional β_1_ and β_2_ AdRs, whose activation by β-AdR agonists such as isoproterenol triggers cAMP-mediated inflammatory responses [25–27]. However, whether such microglial activation influences astrocytic AQP4 polarization remains unknown. Therefore, we hypothesize that microglial activation mediates β-AdR agonist-induced dysregulation of AQP4 polarization in the mouse cortex.

## Materials and method

All experimental protocols were approved by the Institutional Animal Care and Use Committee of Ochanomizu University, Japan (animal study protocols 21005, 22017, 23007, 24017, 25007). All animal experiments were performed according to the guidelines for animal experimentation of Ochanomizu University that conform with the Fundamental Guidelines for Proper Conduct of Animal Experiment and Related Activities in Academic Research Institutions (Ministry of Education, Culture, Sports, Science and Technology, Japan). Efforts were taken to minimize the number of animals used. This study was carried out in compliance with the ARRIVE guidelines.

### Animals

Adult male C57BL/6N mice (aged 8–12 weeks, body weight between 20 g and 30 g) were used for the experiments. Mice were housed in a 12-hour light/dark cycle room, with no more than five mice per cage.

### Craniotomy

After inducing anesthesia with urethane (1.6 g/kg, intraperitoneal injection), the mice were secured in a head fixture. The scalp was then incised, and the periosteum over the target area was removed to expose the skull. A metal plate for the head fixture was affixed to the mouse skull using a glass ionomer resin cement (GC Fuji LUTE BC). Once the cement hardened, Super-Bond (Sun Medical) was applied. Subsequently, an electric drill was used to drill a hole with a diameter of 2 mm above the primary visual cortex. Thereafter, saline was applied for one hour to facilitate recovery.

### Drug administration

Isoproterenol, a beta-adrenergic receptor agonist prepared in saline at a concentration of 200 µM, was dripped into the small hole in the skull and left there for three hours. Minocycline, a microglial antagonist, was continuously injected intraperitoneally for the first two days of the experiment at a dose of 25 mg/kg (Minocycline hydrochloride, CAS No. 13614-98-7, Sigma-Aldrich) for immunostaining and 110 mg/kg for Photothrombosis. Additionally, it was administered once, 20 minutes before the start of the experiment. The PLX diet (containing CSF1R antagonist PLX5622, Chemgood) was administered one week before the experiment, and the control diet (AIN-76A rodent diet, D10001, ResearchDiets) was also administered for one week.

### Immunostaining

Mice were perfused with PBS followed by 4% paraformaldehyde (PFA), and brains were fixed overnight in 4% PFA at 4 °C. The brain was sectioned into 60 μm slices using a micro slicer (DOSAKA, DTK-1000N). Slices were placed in a blocking solution (10 % normal goat serum in PBS with 0.1% TritonX-100) at room temperature and shaken (100 rpm, 1 hour). Primary antibodies were anti-AQP4 (Rabbit, Sigma, A5971, 1:2000), anti-CD31(Rat, BD Biosciences, 550274, 1:1000), anti-GFAP (Proteintech, 16825-1-AP, 1:1000), and anti-Iba1 (Abcam, ab178846, 1:1000). Slices were incubated overnight at 4°C with primary antibodies in blocking solution. After washing 3 times with 0.1% PBST, slices for Iba1 or GFAP immunostaining were incubated with secondary antibody (Alexa Fluor 488, Abcam, ab150077, 1:1000) at room temperature on a shaker (100 rpm) for 1 hour. Slices for AQP4/CD31 immunostaining were incubated with secondary antibodies (Alexa Fluor 488, Abcam, ab150077, 1:1000; Alexa Fluor 594, Invitrogen, A11007, 1:1000) under the same conditions. Afterward, slices were rinsed three times with 0.1% PBST and twice with phosphate buffer (PB) before microscopic examination. Confocal images were acquired using a ZEISS LSM700 microscope and subsequently processed with ImageJ and MATLAB.

### Photothrombosis

Mice were anesthetized with isoflurane (initially 4% for induction, then maintained at 2%), then rose bengal (110 mg/kg; Sigma Aldrich) was injected intraperitoneally and allowed to circulate through the body for 5 minutes. During this period, the mouse was secured in a head holder, and the scalp was incised to expose the skull. A mixture of paraffin oil and Vaseline (1:1) was applied to keep the area moist. A green light (wavelength 532 nm, LED source) was then placed 15 cm from the skull, and the primary visual cortex was irradiated for 15 minutes [15].

### 2, 3, 5-Triphenyltetrazolium staining

Twenty-four hours after photothrombosis, the mice were anesthetized and perfused intracardially with PBS. The excised brains were then placed in 2% 2,3,5-triphenyltetrazolium chloride (T8877, Sigma) in PBS at 40°C for 15 minutes [15].

### Image processing

Fluorescent images captured using a confocal microscope were processed with the Z Project in ImageJ and subsequently analyzed using MATLAB. For images co-staining with AQP4 and CD31, perivascular regions (ROIs) were defined in MATLAB by expanding areas in the CD31 channel that exceeded the fluorescence intensity threshold (mean fluorescence + 1 standard deviation, automatically determined per image) by 3 pixels. The mean fluorescence intensity of AQP4 within these ROIs was then calculated and normalized by the global AQP4 fluorescence intensity [28]. For images of Iba1 and GFAP Immunostaining, the ROIs were determined using MATLAB’s imbinarize function, and the mean fluorescence intensity within each ROI was calculated. Following the method of Young, K., and Morrison, H. (2018), the morphological analysis of microglia was conducted using ImageJ [29]. For all fluorescence images, quantitative measurements were normalized to the left cerebral cortex.

### Statistical analysis

Student’s t-test was used for all the experiments, and all data are expressed as mean ± s.e.m.

## Results

### β-AdR agonist altered the polarization of AQP4

We first hypothesized that β-AdR stimulation affects the polarization of astrocytic AQP4. To test this, we performed cranial window surgery above the primary visual cortex of mice, followed by local administration of a β-AdR agonist for three hours (**Fig. 1A**). Immunostaining revealed that β-AdR agonist administration significantly reducd AQP4 polarization in perivascular regions compared to the saline controls (**Figs. 1B-C**, Saline vs. β-AdR agonist, 94.78 ± 3.07 vs. 76.61 ± 4.83, p = 1.22E-02). These results support our hypothesis that β-AdR stimulation disrupts astrocytic AQP4 polarization.

**Fig. 1.**
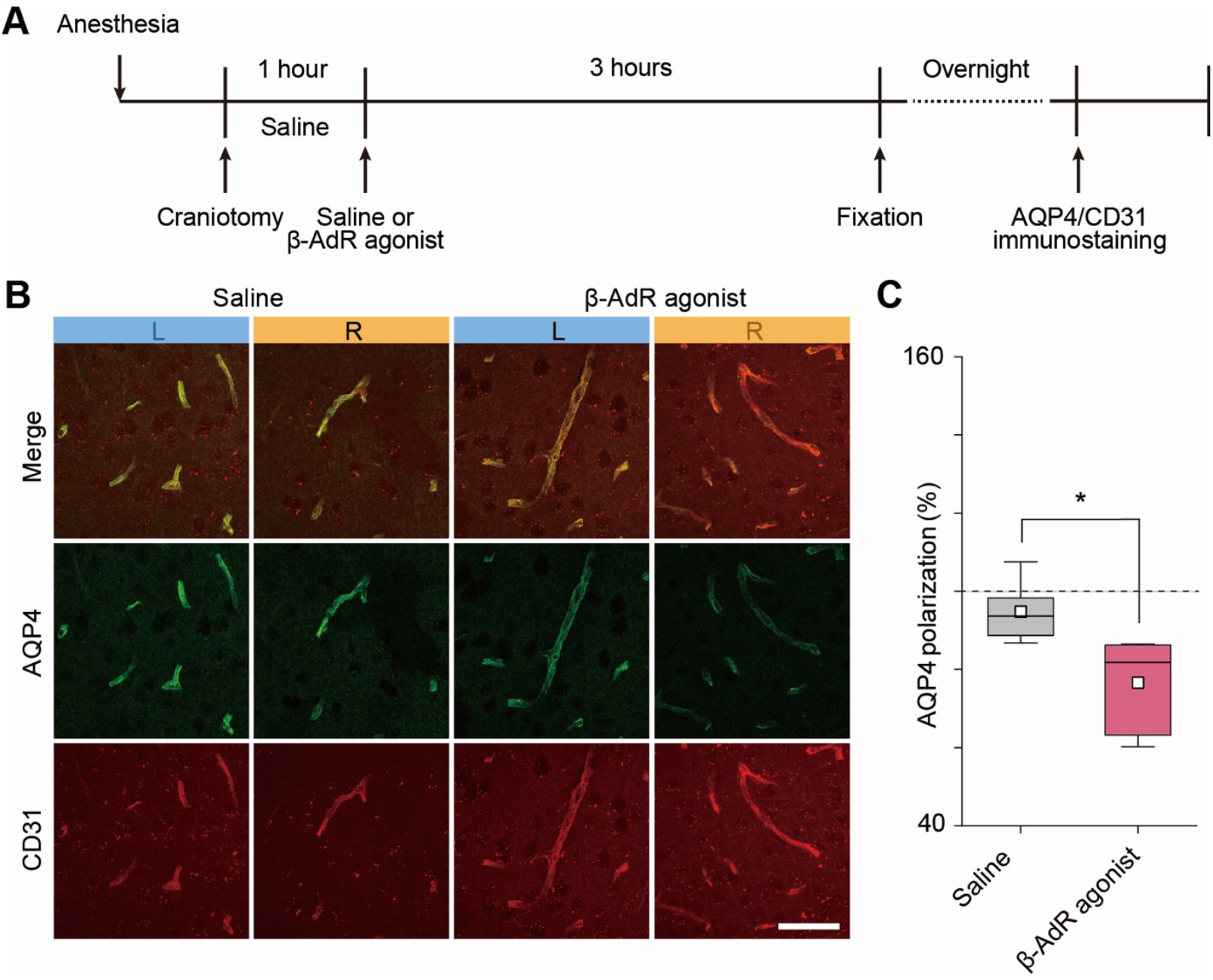
β-AdR agonist alters the polarization of AQP4. (A) Schematic diagram of the experimental protocol. (B) Representative immunostaining images of AQP4 and CD31 from saline and β-AdR agonist-treated mice. Scale bar: 50 μm. Brightness and contrast were uniformly adjusted across images. (C) Quantification of perivascular AQP4 polarization (%) in saline- and β-AdR agonist-treated groups. *p < 0.05 (Saline, n = 6 mice; β-AdR agonist, n = 6 mice, t-test).

### β-AdR agonist enhanced microglial reactivity without affecting astrocytic activation

Microglia express β-adrenergic receptors [25–27], and activated microglia can influence astrocyte function [30,31]. Thus, we hypothesized that microglial activation might mediate the observed changes in AQP4 polarization (**Fig. 2A**). To evaluate this hypothesis, we assessed microglial and astrocytic reactivities using immunostaining for Iba1 and GFAP, respectively (**Figs. 2B-F**). β-AdR agonist treatment did not significantly alter astrocytic GFAP expression (**Fig. 2F**, Saline vs. β-AdR agonist, 101.97 ± 4.22 vs. 104.98 ± 6.39, p = 7.04E-01). However, the treatment significantly increased microglial Iba1 immunoreactivity intensity (**Fig. 2C**, Saline vs. β-AdR agonist, 97.33 ± 3.75 vs. 127.72 ± 10.38, p = 3.15E-02) and microglial density (**Fig. 2D**, Saline vs. β-AdR agonist, 106.63 ± 8.03 vs. 147.76 ± 11.34, p = 1.59E-02). These findings suggest that the β-AdR agonist-induced microglial activation could mediate AQP4 polarization changes.

**Fig. 2.**
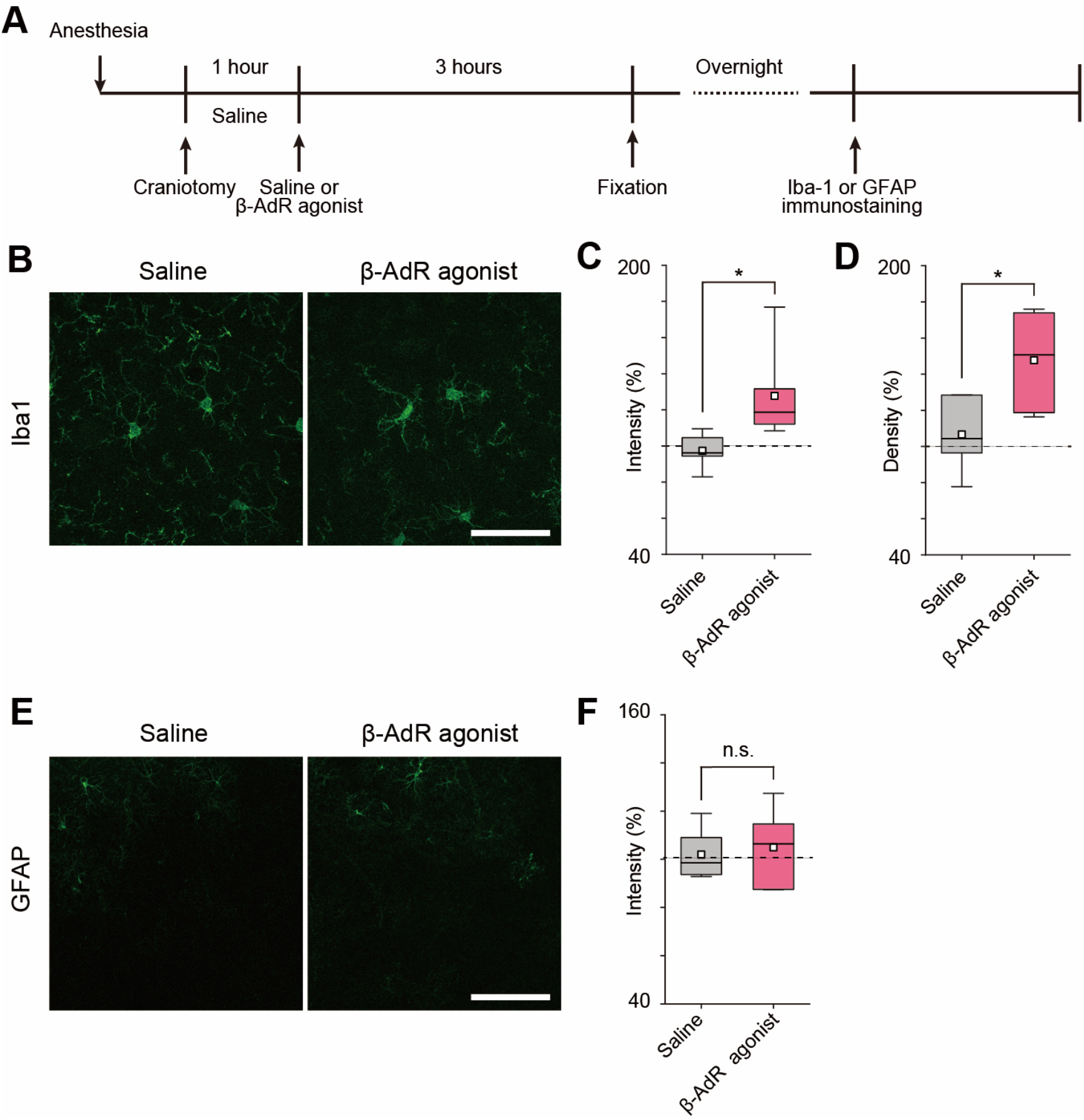
β-AdR agonist alters microglial but not astrocytic activity. (A) Schematic diagram of the experimental protocol. (B) Representative images of Iba1 immunostaining in saline- and β-AdR agonist-treated groups. Scale bar: 50 μm. Brightness and contrast were uniformly adjusted across images. (C) Quantification of Iba1 immunoreactivity intensity (%) in saline- and β-AdR agonist-treated groups. *p < 0.05 (Saline, n = 6 mice; β-AdR agonist, n = 6 mice, t-test). (D) Quantification of microglial density (%) in saline- and β-AdR agonist-treated groups. *p < 0.05 (Saline, n = 6 mice; β-AdR agonist, n = 6 mice, t-test). (E) Representative images of GFAP immunostaining in saline- and β-AdR agonist-treated groups. Scale bar: 100 μm. Brightness and contrast were uniformly adjusted across images. (F) Quantification of GFAP immunoreactivity intensity (%) in saline- and β-AdR agonist-treated groups. n.s., not significant (Saline, n = 6 mice; β-AdR agonist, n = 6 mice, t-test).

### Microglial inhibition and depletion prevented β-AdR-induced AQP4 dysregulation

To directly test the involvement of microglia in β-AdR agonist-induced AQP4 dysregulation, we pharmacologically inhibited microglial activation using minocycline (**Fig. 3A**). After minocycline administration, β-AdR agonist treatment failed to significantly alter microglial reactivity (**Figs. 3B-D**, Iba1 intensity: minocycline + saline vs. minocycline + β-AdR agonist, 96.43 ± 5.26 vs. 108.06 ± 4.75, p = 1.32E-01; Fig. 3F, density of microglia: minocycline + saline vs. minocycline + β-AdR agonist, 112.10 ± 12.71 vs. 114.98 ± 8.95, p = 8.57E-01). Under these conditions, β-AdR agonist treatment also failed to affect AQP4 polarization (**Figs. 3E-F**, minocycline + saline vs. minocycline + β-AdR agonist, 104.04 ± 5.22 vs. 105.60 ± 4.19, p = 8.21E-01).

**Fig. 3.**
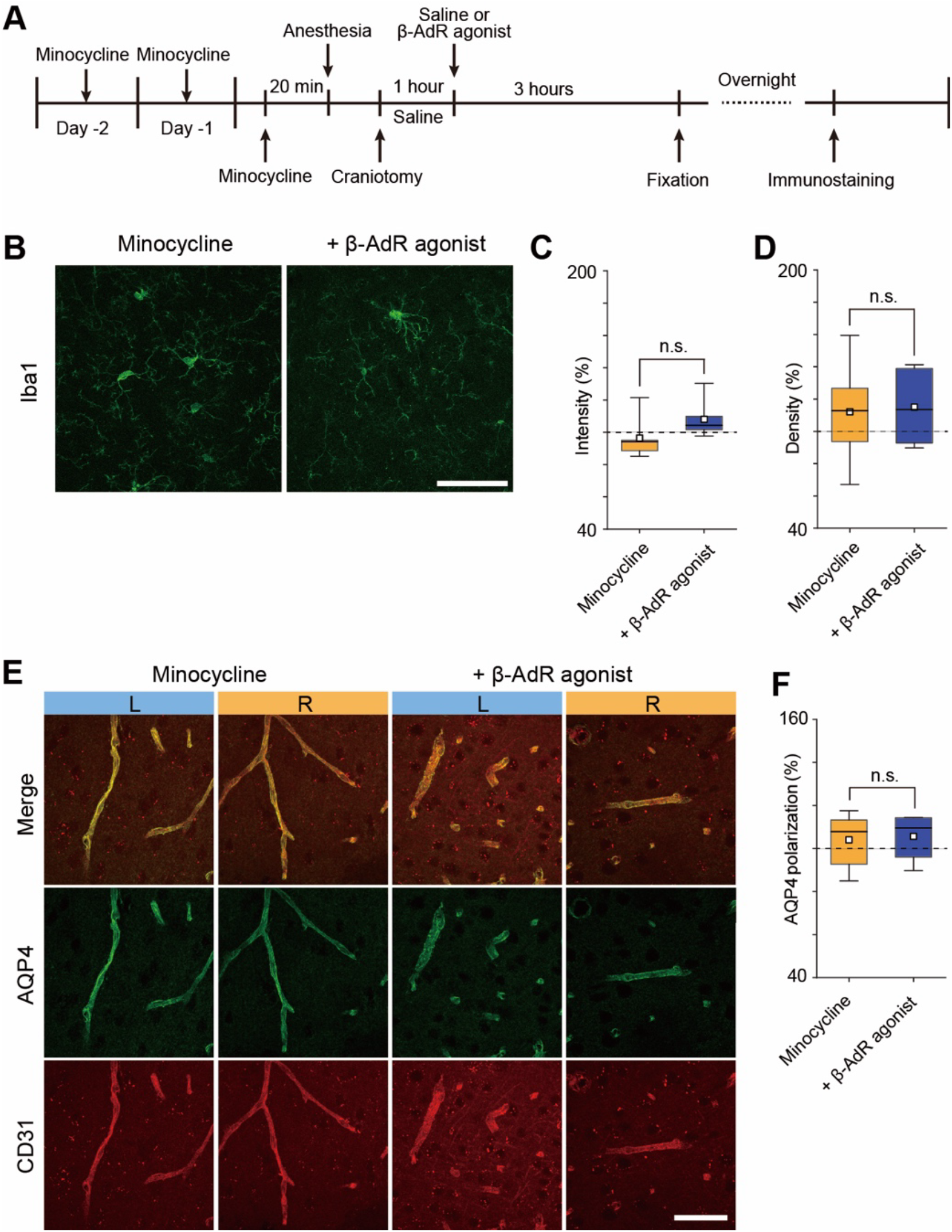
Minocycline prevents β-AdR agonist-induced microglial activation and AQP4 dysregulation. (A) Schematic diagram of the experimental protocol for immunostaining in minocycline + saline and minocycline + β-AdR agonist groups. (B) Representative images of Iba1 immunostaining in minocycline + saline and minocycline + β-AdR agonist groups. Scale bar: 50 μm. Brightness and contrast were uniformly adjusted across images. (C) Quantification of Iba1 immunoreactivity intensity (%) in minocycline + saline and minocycline + β-AdR agonist groups. n.s., not significant (Minocycline + saline, n = 6 mice; Minocycline + β-AdR agonist, n = 6 mice, t-test). (D) Quantification of microglial density (%) in minocycline + saline and minocycline + β-AdR agonist groups. n.s., not significant (Minocycline + saline, n = 6 mice; Minocycline + β-AdR agonist, n = 6 mice, t-test). (E) Representative images of AQP4/CD31 immunostaining in minocycline + saline and minocycline + β-AdR agonist groups. Scale bar: 50 μm. Brightness and contrast were uniformly adjusted across images. (F) Quantification of AQP4 polarization (%) in minocycline + saline and minocycline + β-AdR agonist groups. n.s., not significant (Minocycline + saline, n = 6 mice; Minocycline + β-AdR agonist, n = 6 mice, t-test).

Additionally, we depleted microglia by administering the CSF1R antagonist PLX5622 for one week prior to β-AdR agonist treatment, using the AIN diet as a control (**Fig. 4A**). Although the PLX diet did not significantly alter residual microglial activation following β-AdR agonist administration (**Figs. 4B-D**, Iba1 intensity: AIN + β-AdR agonist vs. PLX + β-AdR agonist, 124.49 ± 6.88 vs. 110.53 ± 12.35, p = 3.14E-01; Fig. 4F, density of microglia: AIN + β-AdR agonist vs. PLX + β-AdR agonist, 134.49 ± 7.43 vs. 118.34 ± 9.86, p = 2.19E-01), β-AdR agonist treatment failed to reduce AQP4 polarization in the microglia-depleted mice (**Figs. 4E-F**, AIN + β-AdR agonist vs. PLX + β-AdR agonist, 85.65 ± 3.35 vs. 102.48 ± 5.34, p = 2.39E-02). These results indicate that microglial activation is essential for β-AdR agonist-induced dysregulation of AQP4 polarization.

**Fig. 4.**
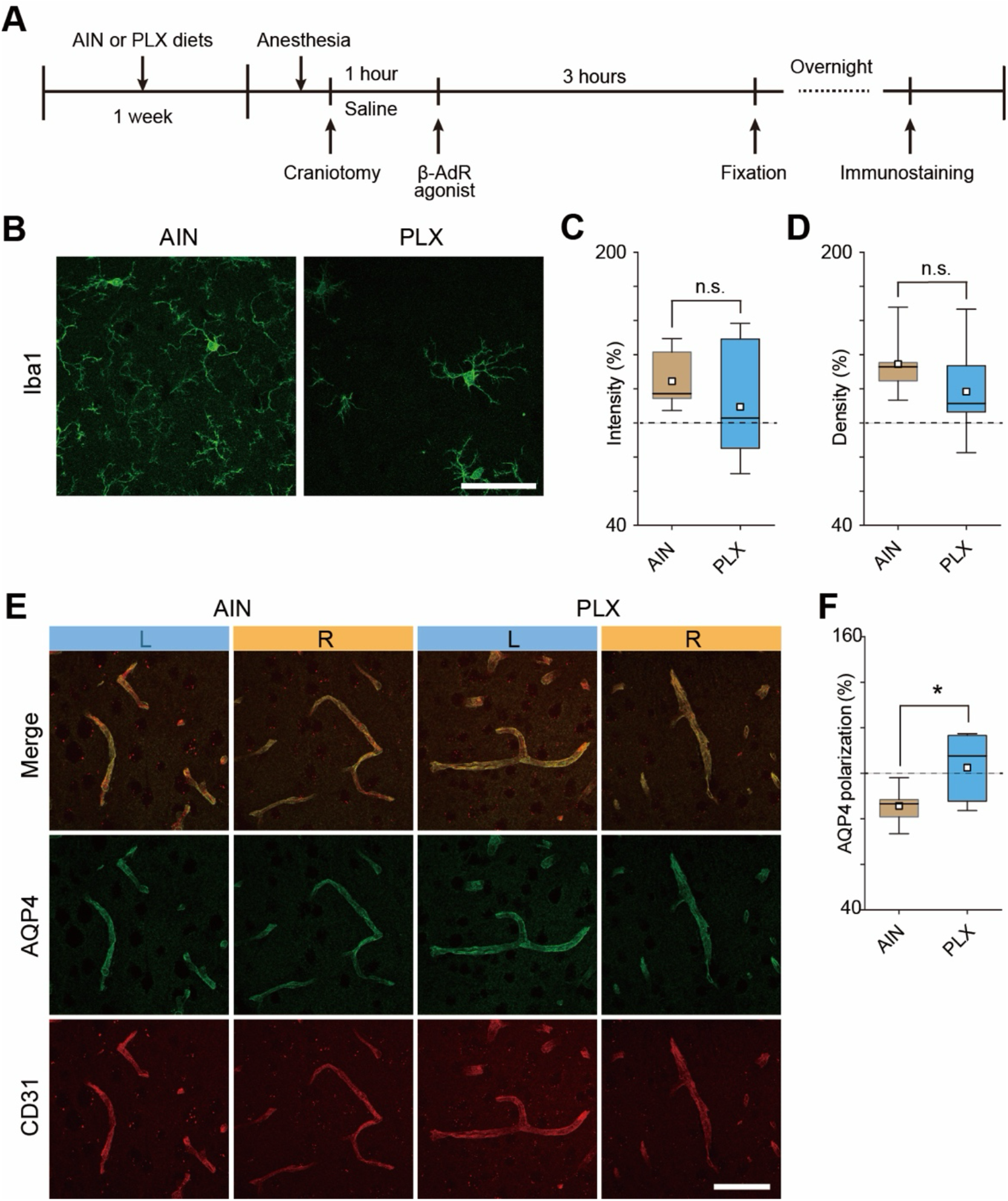
Microglial depletion by PLX5622 prevents β-AdR agonist-induced dysregulation of AQP4 polarization. (A) Schematic diagram of the experimental protocol for immunostaining in AIN + β-AdR agonist and PLX + β-AdR agonist groups. (B) Representative images of Iba1 immunostaining in AIN + β-AdR agonist and PLX + β-AdR agonist groups. Scale bar: 50 μm. Brightness and contrast were uniformly adjusted across images. (C) Quantification of Iba1 immunoreactivity intensity (%) in AIN + β-AdR agonist and PLX + β-AdR agonist groups. n.s., not significant (AIN + β-AdR agonist, n = 6 mice; PLX + β-AdR agonist, n = 7 mice, t-test). (D) Quantification of microglial density (%) in AIN + β-AdR agonist and PLX + β-AdR agonist groups. n.s., not significant (AIN + β-AdR agonist, n = 6 mice; PLX + β-AdR agonist, n = 7 mice, t-test). (E) Representative images of AQP4/CD31 immunostaining in AIN + β-AdR agonist and PLX + β-AdR agonist groups. Scale bar: 50 μm. Brightness and contrast were uniformly adjusted across images. (F) Quantification of AQP4 polarization (%) in AIN + β-AdR agonist and PLX + β-AdR agonist groups. *p < 0.05 (AIN + β-AdR agonist, n = 6 mice; PLX + β-AdR agonist, n = 7 mice, t-test).

### Minocycline reduced infarct area following photothrombosis

Acute cerebral ischemia is associated with microglial activation, which contributes to vasogenic edema [22,32,33]. Loss of perivascular AQP4 polarization further exacerbates to vasogenic edema during acute ischemic injury [34]. To test whether microglial inhibition mitigates ischemic brain injury, we administered minocycline prior to photothrombotic stroke induction (**Fig. 5A**). Minocycline treatment significantly reduced infarct area compared to untreated controls (**Figs. 5B-C**, Untreated vs. Minocycline: 12.80 ± 1.39 mm^2^ vs. 7.91 ± 1.10 mm^2^, p = 2.16E-02). These results indicate that microglial inhibition can ameliorate ischemia-induced brain injury, potentially by preserving AQP4 polarization and subsequently attenuating vasogenic edema.

**Fig. 5.**
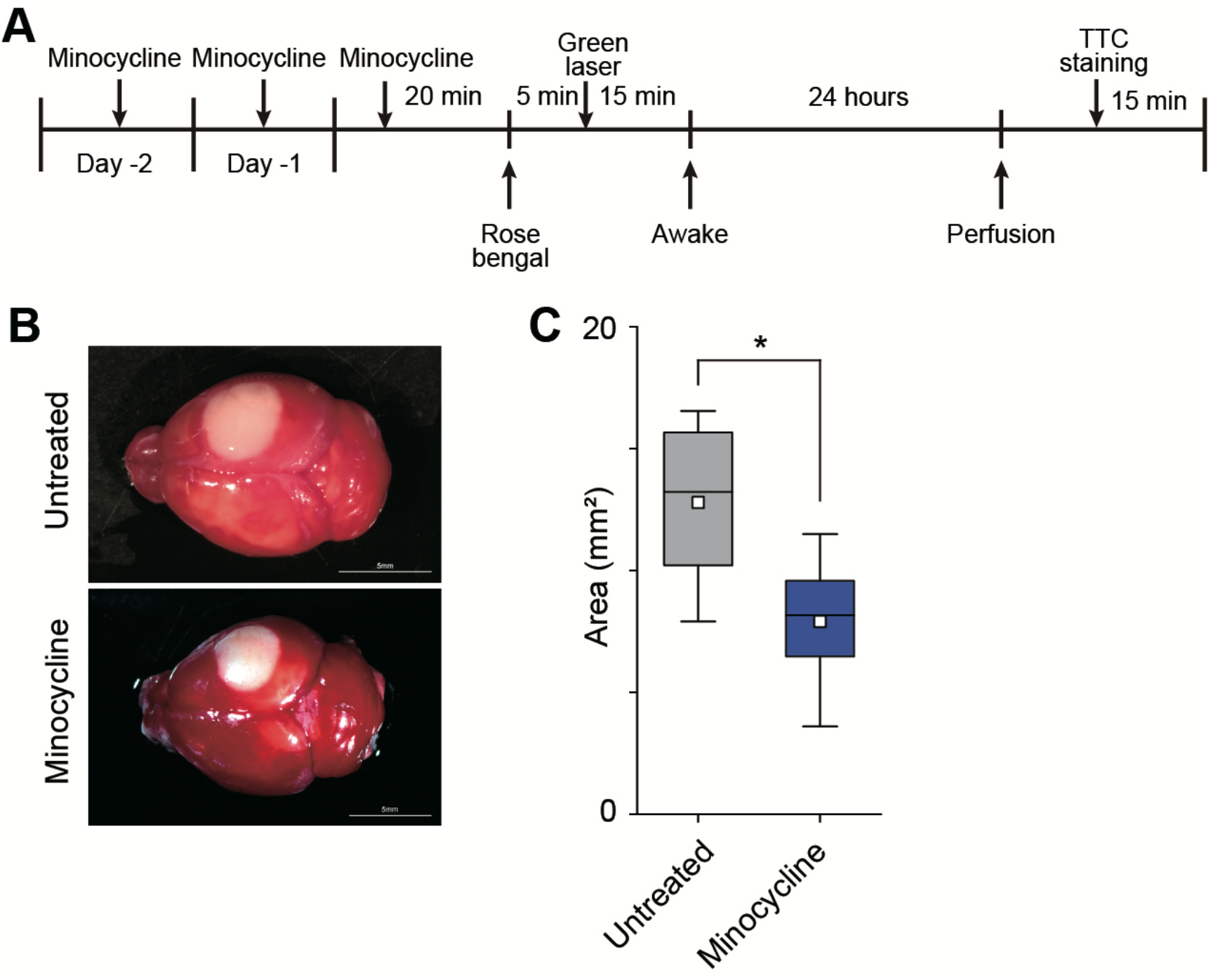
Minocycline reduces infarct area induced by photothrombosis. (A) Schematic diagram of the experimental protocol for evaluating photothrombosis-induced infarction by TTC staining in untreated and minocycline-treated groups. (B) Representative TTC-stained brain sections showing infarct areas in untreated and minocycline-treated groups. (C) Quantitative comparison of infarct area between untreated and minocycline-treated groups. *p < 0.05 (Untreated, n = 6 mice, Minocycline, n = 6 mice, t-test).

## Discussion

In this study, we discovered that β-AdR agonists induce the loss of AQP4 polarization through microglial activation, suggesting that microglia play a crucial role in regulating AQP4 polarization (**Figs. 1-2**). Administration of the β-AdR agonist isoproterenol significantly reduced perivascular AQP4 polarization within three hours (**Fig. 1C**). Since both microglia and astrocytes express β-AdRs [25–27,35], isoproterenol could theoretically activate either cell type. However, astrocytic activation, as assessed by GFAP immunoreactivity, did not change within our experimental time frame (**Fig. 2F**), possibly due to the longer duration required for GFAP protein expression [36]. In contrast, microglia activation, measured by increased Iba1 immunoreactivity, occurred rapidly after β-AdR agonist administration (**Figs. 2C-D**).

To confirm the involvement of microglia, we inhibited their activation pharmacologically with minocycline. Microglial inhibition effectively prevented the β-AdR agonist-induced dysregulation of AQP4 polarization (**Fig. 3F**), supporting our hypothesis that microglial activation mediates this effect. Furthermore, microglial depletion using CSF1R antagonist (PLX5622) also abolished β-AdR agonist-induced dysregulation of AQP4 polarization (**Fig. 4F**), even though residual microglial reactivity markers remained unchanged (**Figs. 4C-D**). Considering that PLX treatment drastically reduces microglial numbers and impairs their function [37], these data suggest that microglial presence and activation status are critical for β-AdR-mediated alterations in astrocytic AQP4 polarization.

Based on these findings, we hypothesize that activated microglia release matrix metalloproteinases (MMPs), particularly MMP-9, [38,39]. MMP-9 cleaves β-dystroglycan (β-DG), a key anchoring protein that stabilizes AQP4 in astrocytic end feet membranes [40,41]. Indeed, activated microglia are a major source of MMP-9 following neuroinflammatory or ischemic insults [23,42]. Thus, the microglia-dependent loss of AQP4 polarization observed here (**Figs. 1-4**) is likely mediated, at least in part, by MMP-9 released from activated microglia, leading to cleavage of β-DG and subsequent mislocalization of AQP4.

Moreover, a recent cellular-level study demonstrated that isoproterenol rapidly reduced AQP4 orthogonal arrays of particles (OAPs), increasing their lateral mobility within astrocytic plasma membranes [43]. Although our study was conducted at the tissue level in vivo (**Fig. 1C, Figs. 3-4**), these findings collectively highlight the complexity of the mechanism by which β-AdR agonists affect AQP4 polarization. Activated microglia promptly release diverse signaling molecules, including ATP [44–47], and astrocytic P2X7 receptor activation by ATP has been shown to downregulate AQP4 expression [48]. Thus, multiple microglia-astrocyte interaction pathways, involving both MMP-mediated proteolytic mechanisms and purinergic signaling pathway, could contribute to the observed depolarization of AQP4.

Finally, several studies suggested that dysregulation of perivascular AQP4 localization exacerbates vasogenic edema in acute cerebral ischemia [14,24,34]. Monai et al. demonstrated that pan-adrenergic receptor antagonists facilitate recovery from acute ischemic stroke by preserving AQP4 polarization [15]. Consistent with these reports, our study shows that minocycline treatment significantly reduced infarct area following photothrombosis-induced ischemia (**Fig. 5**). Minocycline likely exerts its neuroprotective effect by suppressing microglial activation, thereby preventing microglia-dependent disruption of AQP4 polarization and subsequent edema formation.

Taken together, our results reveal a critical role for microglia in adrenergic regulation of astrocytic AQP4 polarization (**Figs. 1-4**) and highlight microglial inhibition as a promising therapeutic strategy for mitigating cerebral edema following acute ischemic injury (**Fig. 5**).

## Acknowledgement

This work was supported by Ochanomizu University, JST FOREST Program, Grant Number JPMJFR204G, Research Foundation for Opto-Science and Technology, Kao Research Council for the Study of Healthcare Science, The Japan Association for Chemical Innovation, and TERUMO LIFE SCIENCE FOUNDATION.

## Notes

### Competing Interest Statement

The authors have declared no competing interest.

### Summary of Updates

The number of animals used has been added to the figure legends.

